# A Kaposi’s Sarcoma-Associated Herpesvirus Infection Mechanism is Independent of Integrins α3β1, αVβ3, and αVβ5

**DOI:** 10.1101/270108

**Authors:** Allison Alwan TerBush, Florianne Hafkamp, Hee Jun Lee, Laurent Coscoy

## Abstract

Host receptor usage by KSHV has been best studied using primary microvascular endothelial and fibroblast cells, although the virus infects a wide variety of cell types in culture and in natural infections. In these two infection models, KSHV adheres to the cell though heparan sulfate (HS) binding, then interacts with a complex of EphA2, xct, and integrins α3β1, αVβ3, αVβ5 to catalyze viral entry. We dissected this receptor complex at the genetic level with CRISPR-Cas9 to precisely determine receptor usage in two epithelial cell lines. Surprisingly, we discovered an infection mechanism that requires HS and EphA2 but is independent of αV- and β1-family integrin expression. Furthermore, infection appears to be independent of the EphA2 intracellular domain. We also demonstrated while two other endogenous Eph receptors were dispensable for KSHV infection, transduced EphA4 and EphA5 significantly enhanced infection of cells lacking EphA2.

**IMPORTANCE:** Our data reveals an integrin-independent route of KSHV infection and suggests that multiple Eph receptors besides EphA2 can promote and regulate infection. Since integrins and Eph receptors are large protein families with diverse expression patterns across cells and tissues, we propose that KSHV may engage with several proteins from both families in different combinations to negotiate successful entry into diverse cell types.

## INTRODUCTION

The rhadinovirus Kaposi’s Sarcoma-Associated Herpesvirus (KSHV) is one of only a handful of human viruses that cause cancers. In the pathogenic state, KSHV infects endothelial cells and leads to the formation of Kaposi’s Sarcoma (KS) tumors which are particularly aggressive in the context of AIDS (1). KSHV infection of B cells can also cause Primary Effusion Lymphoma and Multicentric Castleman’s Disease (2, 3), and the virus has been linked to an inflammatory disorder (KICS) (4, 5). Although treatment with antiretroviral therapy (ART) often leads to resolution of KSHV-related cancers, sudden onset of KS has been recognized as a serious outcome of immune reconstitution inflammatory syndrome (KS-IRIS) following ART (6, 7), and there has been little progress toward controlling KS and KSHV lymphomas in endemic cases.

In the decades since its discovery, it has been observed that KSHV has broad tropism and can efficiently infect many types of human primary cells and cell lines (8–10). KSHV entry mechanisms have been most thoroughly studied in primary microvascular endothelial cells and human foreskin fibroblasts (HFFs), which were of particular interest to understand the origin of the KSHV-infected spindle cells that make up the distinct, highly vascularized KS tumors (reviewed in 11). Infection of monocytes and dendritic cells has also been observed within KS tumors and in tissue culture models (12–14). B cells are thought to be the latently infected reservoir of KSHV (15, 16), but modeling their infection in a laboratory setting has proven to be technically challenging.

However, it is reasonable to assume that the first cells infected in a new host upon transmission are epithelial cells. While KSHV was first considered to be a sexually transmitted infection because of its coinfection pattern with HIV, it is now widely recognized that KSHV can be transmitted through saliva and close contact between individuals (reviewed in 17). Multiple studies have shown that KSHV infects primary human epithelial cells and cell lines including oral keratinocytes (8, 10, 18–22) and another clinical report provides compelling clinical evidence that infection of the tonsillar epithelium could provide a gateway through which the virus might access the underlying lymphocytes to establish the reservoir of latently infected B cells (23).

KSHV interacts with a variety of receptors on the surface of host cells. Heparan sulfate (HS) is thought to be a major cell attachment factor and several KSHV glycoproteins have HS-binding activities (24–30). KSHV also coordinates a complex of integrins α3β1, αVβ3, αVβ5, erythropoietin-producing hepatocellular (Eph) receptor A2 (EphA2), and SLC7A11/xCT to trigger clathrin-mediated endocytosis or macropinocytosis of the virion in HFF cells and primary endothelial cells, respectively (most recently reviewed in 11 and 31). Some questions have been raised over precisely which integrins are required for the infection of individual cell lines (32–35). However, in these two well-characterized infection models, the interaction between KSHV gB and the canonical integrin receptors initiates a signaling cascade of FAK, Src, and PI-3K (36–38 and reviewed in 11). KSHV gH/gL binds EphA2 which amplifies this cascade and coordinates endocytosis effectors together with c-Cbl and myosin IIA (39–44). Still, there are important differences in the entry mechanisms used during infection of HFF and primary microvascular endothelial cells, such as the form of endocytosis used to ultimately internalize the virion, hinting that KSHV initiates different entry processes in different types of cells while using the same receptors.

A smaller number of receptor studies have been performed on a variety of epithelial cell lines, but such a unified model of KSHV receptor usage and entry mechanism has not yet been assembled for any individual cell line. Soluble heparin or enzymatic removal of HS from the cell surface inhibits KSHV infection of human embryonic kidney (HEK) 293 cells and human conjunctival epithelial cells, suggesting that HS is required for epithelial cell infection (24, 32, 45, 46). EphA2 is also clearly important for KSHV infection of several cell lines. Soluble EphA2 or Eph-blocking ligands inhibit infection of HEK293 and SLK cells, and EphA2 becomes phosphorylated upon infection in these two cell lines (39, 47–49). Furthermore, siRNA knock down of EphA2 significantly reduces infection of SLK cells (39, 48). Soluble EphA2 inhibits infection of two additional epithelial cell lines (HeLa and HepG2), and overexpression of EphA2 enhances infection of HEK293 cells and the human lung epithelial cell line H1299 (39).

The evidence for integrin involvement during infection of epithelial cell lines is mixed. Two groups have reported that integrin ligands, RGD peptides, soluble α3β1, or function-blocking integrin αV and β1 antibodies did not block KSHV infection of a HEK293-derived reporter cell line or HEK293 cells (32, 50). A third group reported that soluble integrins α3β1 and αVβ3 and a function-blocking αVβ3 antibody did not block KSHV infection of SLK cells (39). However, a fourth group reported that soluble integrins α3β1, αVβ3, and αVβ5 reduced the infection rate of HEK293 cells and that the signaling proteins FAK, ERK1/2, and RhoA were activated upon KSHV infection (37, 51, 45). Finally, a fifth group studied a HeLa-derivative cell line misidentified as human salivary gland epithelial cells HSG(HeLa) and reported that the cells were resistant to KSHV despite expressing all known receptors except integrin β3, and expression of integrin β3 (and restoration of integrin αVβ3) greatly increased the susceptibility of the cells to KSHV infection (35, 52).

Here, we used CRISPR-Cas9 as a tool to comprehensively examine the use of this KSHV receptor complex in two highly infectible epithelial cell lines: Caki-1 kidney epithelial cells, and HeLa cervical epithelial cells. Caki-1 cells are significant as they have contaminated all known stocks of the SLK cell line used in KSHV research (53). We found that HS and EphA2 were required for infection of both Caki-1 and HeLa cells, while αV- and β1-family integrins were dispensable. Interestingly, we also found that the intracellular domain of EphA2 was not required for infection of these cells. Moreover, the ectopic expression of EphA5 and overexpression of EphA4 promoted infection in *EPHA2* knock out (KO) cells but knock out of endogenous EphA4 lead to an elevated infection rate in both WT and *EPHA2* KO contexts. Finally, we also found that infection of primary gingival keratinocytes (PGKs) was unaffected by integrin- or Eph-blocking reagents. Together with other recent studies, our results point to the existence of another unknown KSHV receptor which could trigger intracellular signaling and virion internalization in all three of the cell types we investigated. Our studies revealed a novel KSHV infection mechanism in Caki-1 and HeLa cells that is independent of integrins α3β1, αVβ3, and αVβ5 and suggest that Eph receptors may play more diverse and complex roles during infection than was previously known.

## RESULTS

### Caki-1 and HeLa cells express most known KSHV receptors

It has been shown that KSHV uses a multimolecular complex of attachment molecules and receptors, including HS, EphA2, xCT, DC-SIGN (in some immune cells), and the integrin heterodimers α3β1, αVβ3, and αVβ5, to enter cells in several different infection models (reviewed in 11). The expression of these known KSHV receptors on the surface of Caki-1 and HeLa cells was examined by flow cytometry. Most of the KSHV receptors were expressed on the surface of both cell lines: EphA2, HS, and integrin subunits α3, αV, β1, and β5 (summarized in Fig. 1 and in detail in Supp. Fig. 1). Integrin β3 was additionally detected on the surface of Caki-1 cells but not HeLa cells (Fig. 1, Supp. Fig. 1). However, neither the myeloid cell marker DC-SIGN nor xCT were detected on the surface of either cell line (Fig. 1, Supp. Fig.1).

**Figure 1.**
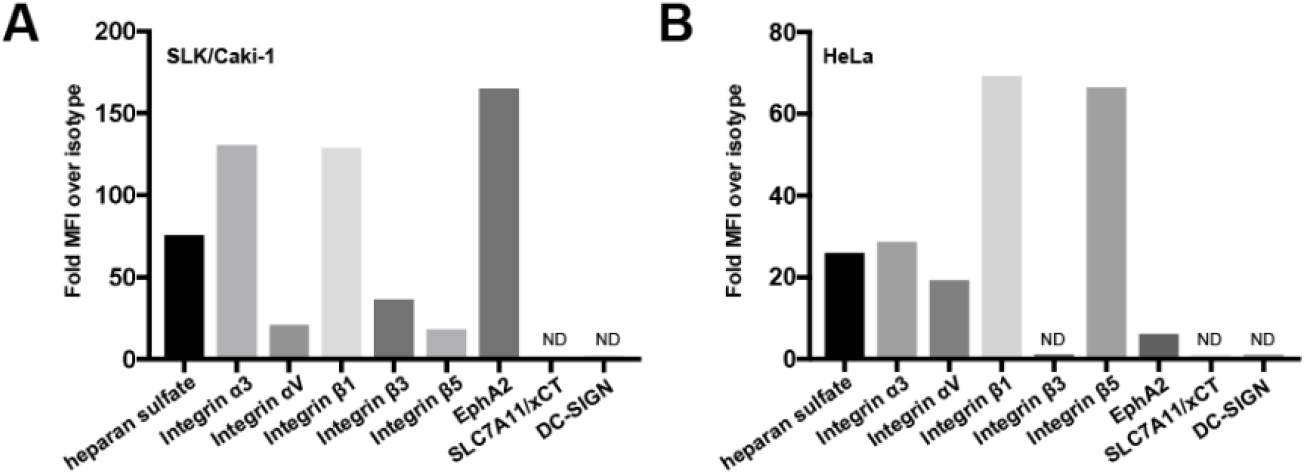
Surface expression of known KSHV receptors on Caki-1 and HeLa cells. Live Caki-1 (A) and HeLa (B) cells were immunostained for surface expression of known KSHV receptors and analyzed by flow cytometry. The mean fluorescence intensity (MFI) of each receptor stain was divided by that of the appropriate primary antibody isotype control. ND, not detected.

### Heparan sulfate interactions are required for KSHV infection of Caki-1 and HeLa cells

The role of HS in adhering virions to the cell surface and promoting viral entry is well documented across many virus families. Caki-1 and HeLa cells express HS on the cell surface and we expected this proteoglycan to play a major role during KSHV infection. We have previously shown that a deficiency in the enzyme Ext1 rendered cells unable to synthesize HS (55), so we could use *EXT1* KO cells to confirm the requirement for HS during KSHV entry.

An *EXT1*-specific guide sequence was cloned into px330, a Cas9 and sgRNA delivery plasmid, which was then transfected into Caki-1 cells (Table 1). After four days, a subpopulation of HS-low mutant cells was discernable by flow cytometry. The mutant population was enriched by fluorescence-activated cell sorting (FACS), then passaged until the immunostaining of HS in the pool decayed to isotype levels (Fig. 2A). It should be noted that this *EXT1* KO pool is polyclonal in nature, and presumably contains cells derived from a multitude of individual CRISPR-Cas9 editing events. This approach helps mitigate the chance of off-target effects contributing significantly to any effects on infection.

**Table 1.**
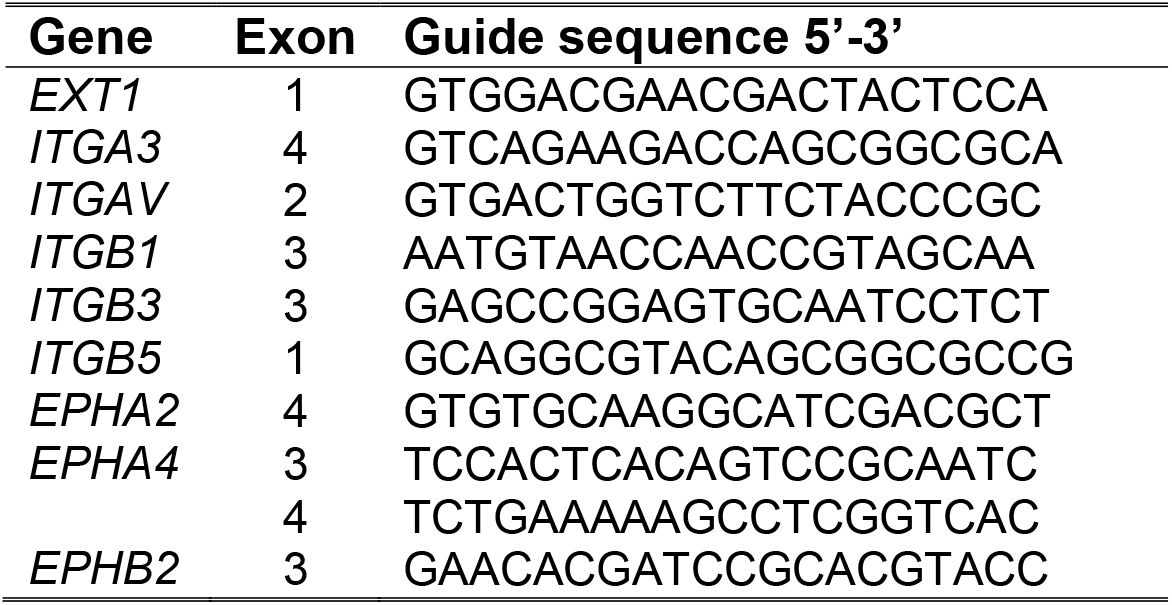
CRISPR-Cas9 guide RNA sequences used to target the indicated genes.

**Figure 2.**
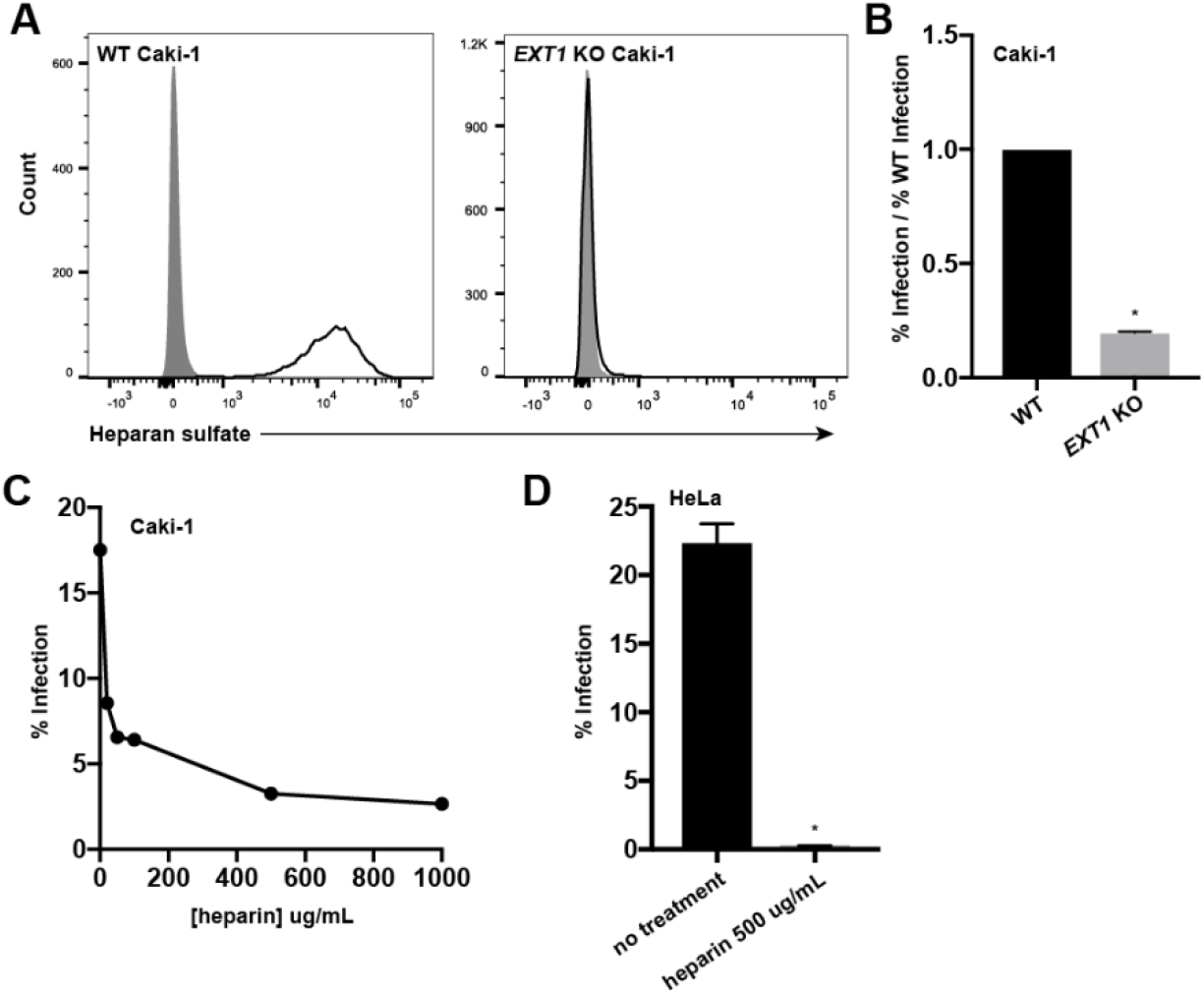
Heparan sulfate interactions are required for infection of Caki-1 and HeLa cells. (A) WT and *EXT1* KO Caki-1 cells were immunostained for surface heparan sulfate (HS) expression. Grey histograms represent isotype controls. (B) WT and *EXT1* KO Caki-1 cells were infected with KSHV in duplicate and infection rates were measured by flow cytometry. The infection rate of the KO was normalized to the average WT infection rate and data was pooled from multiple experiments. (C) Filtered KSHV was pre-incubated with the indicated concentrations of soluble heparin at 37°C then used to infect Caki-1 cells for two hours at 37°C. Infection percentages were measured by flow cytometry two days post infection. (D) Filtered KSHV was pre-blocked with 500 μg/mL of heparin at 37°C, then used to infecte WT HeLa cells in triplicate for two hours at 37°C. Infection percentage was measured by flow cytometry two days post infection. *, p < 0.05.

The *EXT1* KO Caki-1 pool and WT Caki-1 cells were infected with KSHV.BAC16 which encodes a constitutive GFP reporter (56), and the infection percentage was quantified by measuring GFP+ cells by flow cytometry after two days. As expected, the infection rate of HS-deficient Caki-1 cells was significantly reduced compared to WT cells (Figs. 2B).

As an orthogonal approach, we used soluble heparin to competitively block KSHV infection, since this method has been used extensively to investigate HS usage in a variety of cell types (24, 27, 57, 58). KSHV was pre-incubated with increasing concentrations of heparin and then used to infect Caki-1 cells. In agreement with existing literature and our results with *EXT1* KO Caki-1 cells, soluble heparin inhibited infection in a dose-dependent manner but approached a non-zero asymptote (Fig. 2C). Pre-blocking KSHV with 500 μg/mL of heparin had an even more dramatic effect on HeLa cells, completely abrogating infection compared to non-treated virus (Fig. 2D). Collectively, these results show that HS interactions are required for infection of both Caki-1 and HeLa cells and demonstrate the value of CRISPR-Cas9 to study viral receptors.

### KSHV infection of Caki-1 and HeLa cells is independent of canonical KSHV integrin receptors

KSHV coordinates several integrin heterodimers to initiate signaling events that are required for infection of HFFs and primary microvascular endothelial cells (reviewed in 11). Because we observed the expression of all proposed integrin receptors for KSHV at the surface of Caki-1 cells and all, except integrin β3, on the surface of HeLa cells (Fig. 1, Supp. Fig. 1), we investigated whether these integrins were required for KSHV to infect these cell lines. Both the α and β subunits contribute to the unique ligand-binding surface of a given integrin heterodimer. Therefore, we reasoned that infecting cells with reciprocal subunits of KSHV-associated integrins knocked out would reveal precisely which heterodimers were required for infection.

Single KO pools of integrins α3, αV, β1, β3, and β5 were created by transfection of Caki-1 cells with px330 plasmids containing guide sequences that targeted the genes encoding each integrin subunit (Table 1). The mutant populations were enriched as described for *EXT1* to generate integrin KO Caki-1 cell pools (Fig. 3A). Lacking integrin αV protein, *ITGAV* KO cells lost the ability to adhere to tissue-culture treated polystyrene dishes, but normal morphology and growth returned when they were plated on fibronectin-coated plates. The *ITGAV* KO cells were grown on fibronectin for passaging and infection experiments. The infection percentages of both WT Caki-1 and HeLa cells were not affected by the presence of a fibronectin plate coat (data not shown). WT HeLa cells were additionally transfected with px330 plasmids targeting *ITGAV* or *ITGB1*, but the transfected populations were not enriched. The cells were passaged until the receptor expression of the mutant population decayed to near-isotype levels, generating mixed WT/ KO HeLa pools (Fig. 3E). The mixed *ITGAV* KO HeLa pool was also grown on fibronectin.

**Figure 3.**
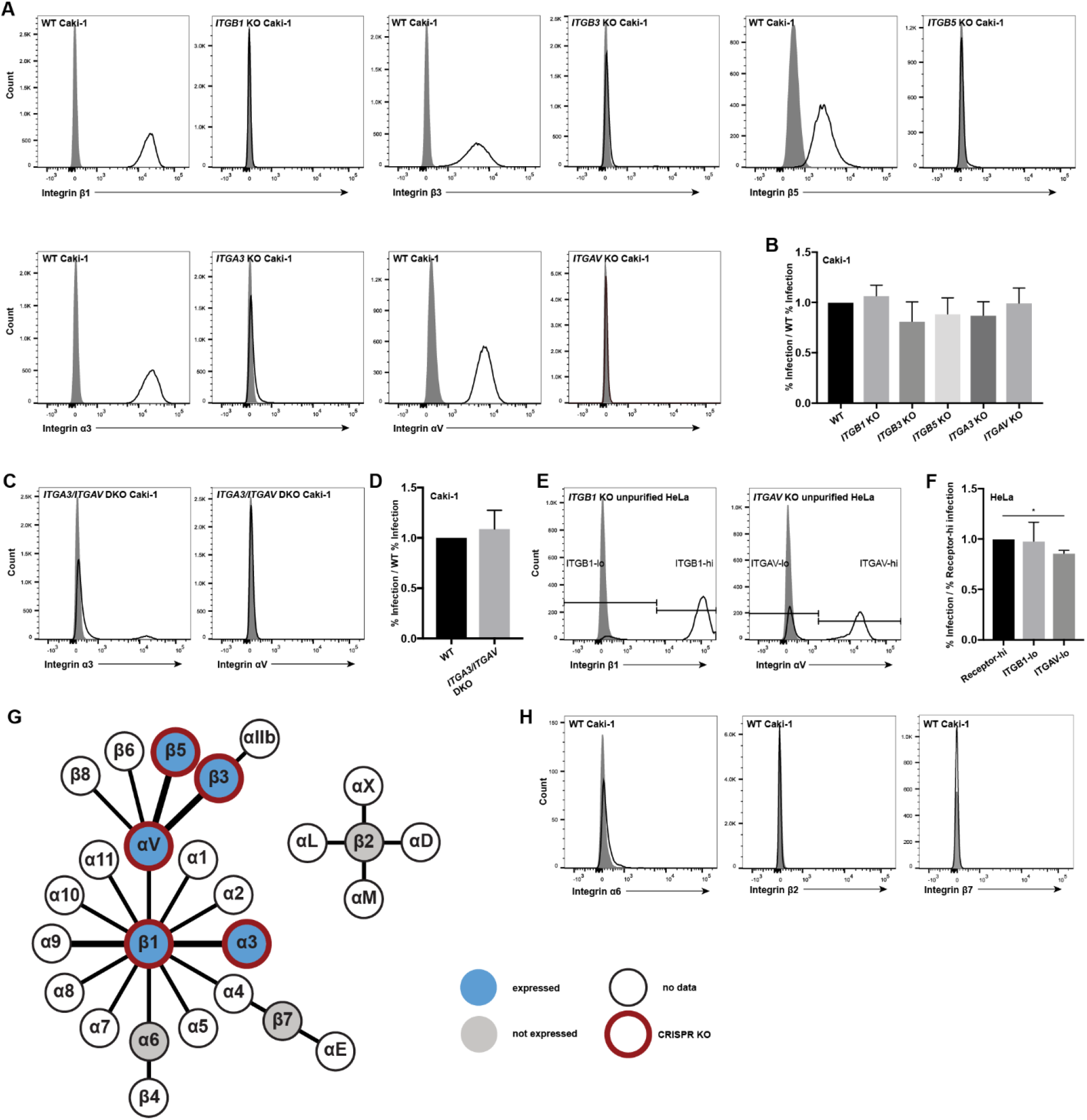
Canonical KSHV integrin receptors are not required for infection of Caki-1 and HeLa cells. (A, H) WT and indicated integrin subunit KO Caki-1 cells were immunostained for surface expression of the indicated integrins. Grey histograms represent isotype controls. (B) WT and integrin KO Caki-1 pools were infected with KSHV in duplicate and infection percentage was measured by flow cytometry two days post infection. The infection rates of the KO pools were normalized to the average WT infection rate and data was pooled from multiple experiments. (C) *ITGA3/ITGAV* double KO Caki-1 cells were immunostained for surface integrin α3 and αV expression. Grey histograms represent the isotype controls. (D) WT and *ITGA3/ITGAV* double KO Caki-1 cells were infected with KSHV in triplicate and infection rate was quantified by flow cytometry. The infection rate of the DKO pool was normalized to the average WT infection rate and data was pooled from multiple experiments. (E) Mixed *WT*/*ITGAV* KO and *WT/ITGB1* KO HeLa populations were immunostained for surface integrin αV and β1 expression. Grey histograms represent the isotype controls. (F) Mixed integrin KO HeLa pools were infected with KSHV in triplicate and infection percentages were measured by flow cytometry two days post infection. The pools were also immunostained for the corresponding integrins and gated on integrin-high or -low populations as indicated in (E). The infection rates of integrin-low cells were normalized to integrin-high cells in each well and a representative experiment is shown. *, p < 0.05. (G) Schematic integrin pairing diagram (adapted from 79) showing expression data measured by surface immunostaining and flow cytometry and subunits targeted by our CRISPR-Cas9 KO approach. Bold connections denote heterodimers previously implicated in KSHV infection.

WT Caki-1, the single integrin KO Caki-1 pools, and the mixed integrin KO HeLa cell pools were then infected with KSHV. The mixed KO HeLa pools were additionally stained for surface expression of the appropriate integrin at the time of infection analysis to allow for gating on WT and KO subpopulations. Overall, the infection percentages of the integrin KO pools or subpopulations were not significantly reduced compared to that of WT cells for both cell lines (Fig. 3B, 3F). The slight decline in infection percentage of the *ITGAV* KO HeLa subpopulation compared to the WT subpopulation reached statistical significance, but the magnitude of difference was small and similar to the other integrin KO Caki-1 pools. Since the KO pools were enriched by FACS, it is likely that there were still a small number of cells that expressed WT levels of each integrin subunit. Nevertheless, these data suggest that KSHV infection of Caki-1 and HeLa cells does not require integrin α3β1, αVβ3, or αVβ5 alone, or any other single integrin in the αV and β1 families.

Although targeting single integrin heterodimers has yielded clear infection defects in past studies (34–36, 45), we considered that our strategy of knocking out individual integrin subunits would not reveal fully redundant involvement of α3β1, αVβ3, and αVβ5 during infection of Caki-1 cells. To address this, an *ITGA3/ITGAV* double KO Caki-1 pool was generated to effectively deplete integrin α3β1 and the entire integrin αV family, including αVβ3 and αVβ5, from the cell surface (Fig. 3C). As with previous receptor KO pools, the *ITGA3/ITGAV* DKO pool was enriched but not purified, since a very small population of cells expressing WT levels of integrin α3 was still visible by flow cytometry (Fig. 3C). WT and *ITGA3/ITGAV* double KO Caki-1 cells were then infected with KSHV. Still, the infection percentage of *ITGA3/ITGAV* double KO Caki-1 cells was not significantly reduced compared to WT cells at either one or two days post infection (Fig. 3D, Supp. Fig. 5). These results further indicate that integrins α3β1, αVβ3, and αVβ5 are not required for KSHV infection of Caki-1 cells.

We also considered whether our genetic disruptions in the integrin network were altering the expression level of other KSHV receptors, potentially obscuring an infection defect in integrin subunit KO cells. To address this, we examined the expression of all known KSHV receptors on *ITGB1* KO and *ITGAV/ITGA3* DKO Caki-1 cells (Supp. Fig. 2). We observed that *ITGB1* KO cells lost surface expression of integrin α3, which is not unexpected since integrin α3 does not bind to any other known integrin β subunits (Supp. Fig. 2). Likewise, we found that *ITGAV/ITGA3* DKO Caki-1 cells lost surface expression of integrins β3 and β5 (Supp. Fig. 2). Otherwise, we did not observe any large changes in the surface expression of unrelated integrin subunits, HS, or EphA2 (Supp. Fig. 2).

Past studies have utilized integrin-blocking reagents to show that certain classes of integrins are required for KSHV entry in a variety of cell types (34, 36, 45, 59). However, at least three publications have reported that several integrin-blocking reagents failed to inhibit KSHV infection in HEK239 and SLK cells (32, 39, 50). To confirm that our results were not unique to the CRISPR-Cas9 KO approach, we replicated key integrin-blocking experiments from these publications. WT Caki-1 and HeLa cells were pre-incubated with the RGD-containing integrin ligand fibronectin, the integrin ligand laminin, GRGDSP and GRGESP peptides, or a DMSO control for the peptide resuspension solution for one hour at 4°C. The cells were then washed and infected with KSHV for two hours at 37°C. Fibronectin, which contains an RGD sequence and binds αV-family integrins, did not significantly alter infection rate of either cell line (Supp. Fig. 3). Laminin, which binds to a different subset of integrins including α3β1, slightly inhibited infection of HeLa cells but not Caki-1 cells (Supp. Fig. 3). Neither the RGD-containing peptide GRGDSP nor the control peptide GRGESP significantly affected KSHV infection of HeLa cells (Supp. Fig. 3). GRGDSP very slightly inhibited infection of Caki-1 cells, but the effect was not significantly different compared to GRGESP, suggesting that the inhibitory effect of the peptide was nonspecific which has been noted in a previous publication (32) (Supp. Fig. 3). Overall, we found that these blocking reagents had little or no effect on KSHV infection in Caki-1 and HeLa cells which was consistent with the results of our CRISPR-Cas9 KO studies.

A non-RGD-binding integrin, α9β1, has been shown to bind a disintegrin-like domain (DLD) in KSHV gB and is important for infection of HFF and primary microvascular endothelial cells, but not HEK293 cells (50). Our data demonstrate that KSHV infection of Caki-1 and HeLa cells is independent of the twelve β1-containing integrins and the five αV-containing integrins, however we considered that other integrins could still be required for KSHV infection of Caki-1 cells. There are eight integrins that do not contain the αV or β1 subunits: αIIbβ3, α6β4, α4β7, αEβ7, and four β2-containing integrins (Fig. 3G). Neither integrin α6, integrin β7, nor integrin β2 were detected on the surface of WT Caki-1 cells by flow cytometry (Fig. 3H). Additionally, integrin β3 was lost from the cell surface of *ITGAV/ITGA3* DKO Caki-1 cells implying that integrin αIIbβ3 is not expressed in Caki-1 cells (Supp. Fig. 2). Altogether these data indicate that none of the eight non-αV, non-β1 integrin heterodimers are expressed in Caki-1 cells, so these integrins are unlikely to play a role in this KSHV infection mechanism in the absence of αV- or β1-family integrins.

### EphA2 is necessary for KSHV infection of Caki-1 and HeLa cells

EphA2 has been well-characterized as a receptor for KSHV and binds to the envelope glycoprotein complex gH/gL (39, 40, 44, 49). Together with integrins, EphA2 helps propagate virus-induced signaling and mobilize endocytosis effectors which leads to viral entry in multiple cell types (39–43, 48). However, we found that KSHV infection of Caki-1 and HeLa cells does not require canonical KSHV integrin receptors, so we investigated whether EphA2 was required for infection in these cell lines.

Caki-1 cells were transfected with a px330 plasmid containing a guide sequence targeting *EPHA2* and an *EPHA2* KO pool was enriched as described for *EXT1* (Table 1, Fig. 4A). In addition, a mixed *WT*/*EPHA2* KO pool was generated in HeLa cells as described for *ITGAV* and *ITGB1* (Fig. 4D). WT and *EPHA2* KO Caki-1 cells and the mixed WT/KO HeLa pool were then infected with KSHV. The mixed WT/KO HeLa pool was additionally stained for surface EphA2 expression at the time of infection analysis to distinguish the KO and WT subpopulations. The infection percentage of *EPHA2* KO cells was significantly reduced compared to WT cells in both cell lines, though *EPHA2* KO cells were not completely resistant to the virus (Figs. 4B, 4E). The infection defect in *EPHA2* KO cells was similar at both one and two days post infection (Supp. Fig. 5). These results indicate that EphA2 is necessary for infection of both Caki-1 and HeLa cells.

**Figure 4.**
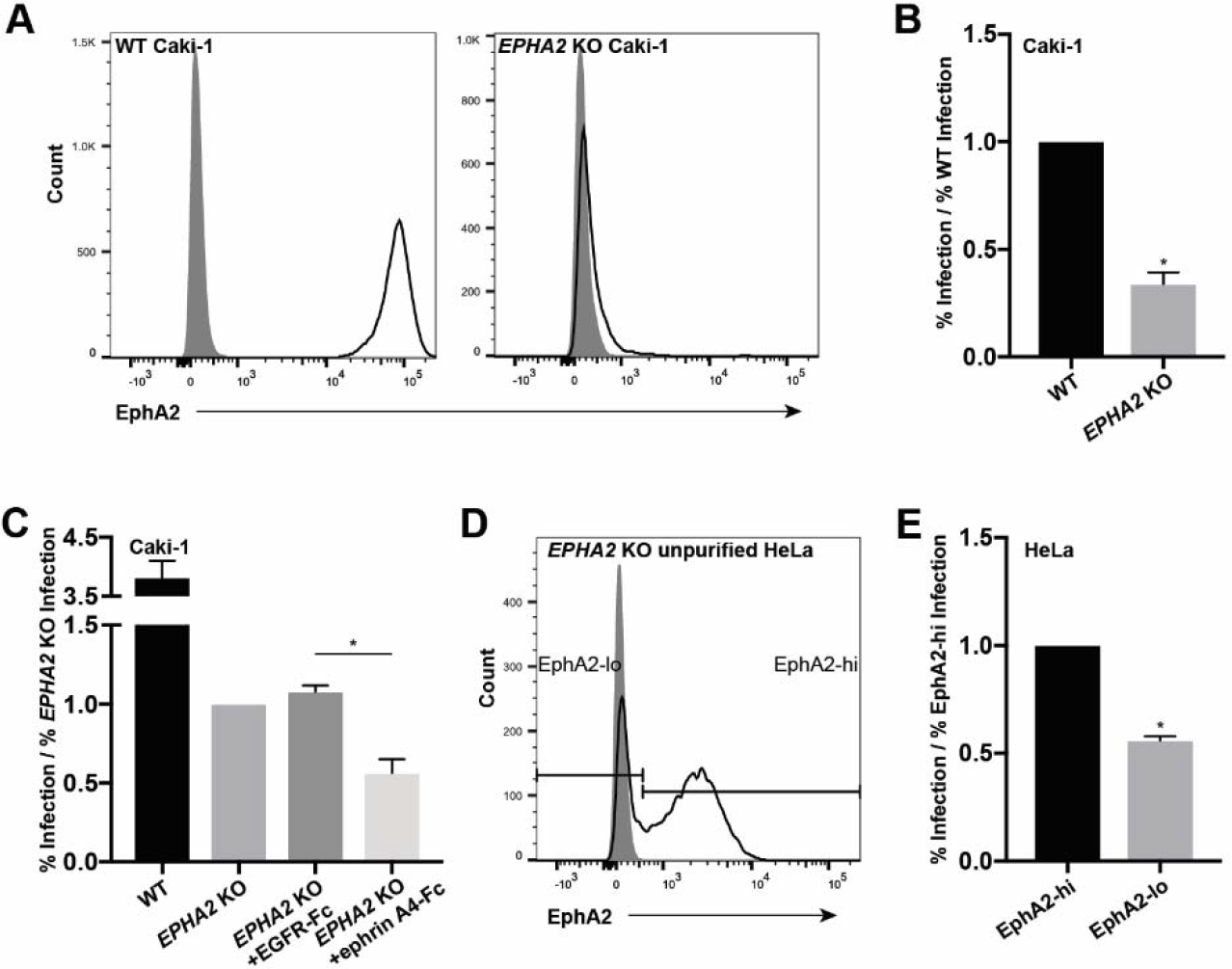
EphA2 is required for infection of Caki-1 and HeLa cells. (A) WT and *EPHA2* KO Caki-1 cells were immunostained for surface EphA2 expression with the SHM16 antibody and analyzed by flow cytometry. Grey histogram represents isotype control. (B) WT and *EPHA2* KO Caki-1 cells were infected with KSHV in duplicate and infection rates were quantified by flow cytometry. The infection rate of the *EPHA2* KO pool was normalized to the average WT infection rate and data was pooled from multiple experiments. (C) *EPHA2* KO Caki-1 cells were pre-blocked with EGFR- Fc or ephrin-A4-Fc at 10 μg/mL at 4°C and then infected in triplicate in the presence of EGFR-Fc or ephrin-A4-Fc at 5 μg/mL at 37°C. Infection percentage was measured by flow cytometry two days post infection and percent infection was normalized to the average *EPHA2* KO infection rate. (D) Mixed *EPHA2* KO HeLa cells were immunostained for surface EphA2 expression with the SHM16 antibody and analyzed by flow cytometry. Grey histogram represents the isotype control. (E) The mixed *EPHA2* KO HeLa cells were infected with KSHV in triplicate and infection percentage was measured by flow cytometry two days post infection. The cells were also immunostained for surface EphA2 and gated on EphA2-high or -low as indicated in (D). The infection rates of EphA2-low cells were normalized to EphA2-high cells in each well and a representative experiment is shown. *, p < 0.05.

To ensure that the KO of *EPHA2* did not alter the expression of any other known KSHV receptors, WT and *EPHA2* KO Caki-1 cells were examined for surface receptor expression by flow cytometry. We did not observe any unexpected changes in the surface expression of any other known receptors (Supp. Fig. 2).

A prior study demonstrated that KSHV gH/gL has the ability to bind other Eph receptors besides EphA2 *in vitro* (47). We hypothesized that EphA2-independent infection could depend on other Eph receptors that may be expressed. A clonal EphA2 KO Caki-1 cell line was isolated from single-cell clones of the EphA2 KO pool and was used for this experiment. The clone lacked surface expression of EphA2 and the infection defect compared to WT cells was similar to the parent population (Figs. 4C, 6E). To ensure this EphA2 KO clone did not produce EphA2 protein, surface and total EphA2 were examined by flow cytometry and western blot, respectively, using a second EphA2-specific antibody (Supp. Fig. 4). To test whether additional Eph receptors were required for infection in these cells, we attempted to block KSHV infection using the A-type Eph ligand ephrin-A4 or EGFR as a control, as reported previously (39, 47). Clonal *EPHA2* KO cells were pre-incubated with either soluble ephrin-A4-Fc or EGFR-Fc chimeric proteins, then infected with KSHV in the presence of these blocking agents. The infection percentage of *EPHA2* KO cells was further reduced in the presence of ephrin-A4-Fc compared to the unrelated EGFR-Fc (Fig. 4C). Since ephrin ligands, including ephrin-A4, can broadly bind to and block interactions with Eph receptors of the same type, these data suggest that another A-type Eph receptor may be required for infection of Caki-1 cells in the absence of EphA2.

### EphA4 and EphB2 are dispensable for KSHV infection in Caki-1 cells

Since we found that KSHV infection in *EPHA2* KO Caki-1 cells could be further blocked by a soluble ephrin ligand, we investigated whether additional Eph receptors were expressed by Caki-1 cells and if they were required for KSHV infection of Caki-1 cells. EphA4 and EphB2 were found to be expressed by Caki-1 cells by western blot (Figs. 5E, 5H).

**Figure 5.**
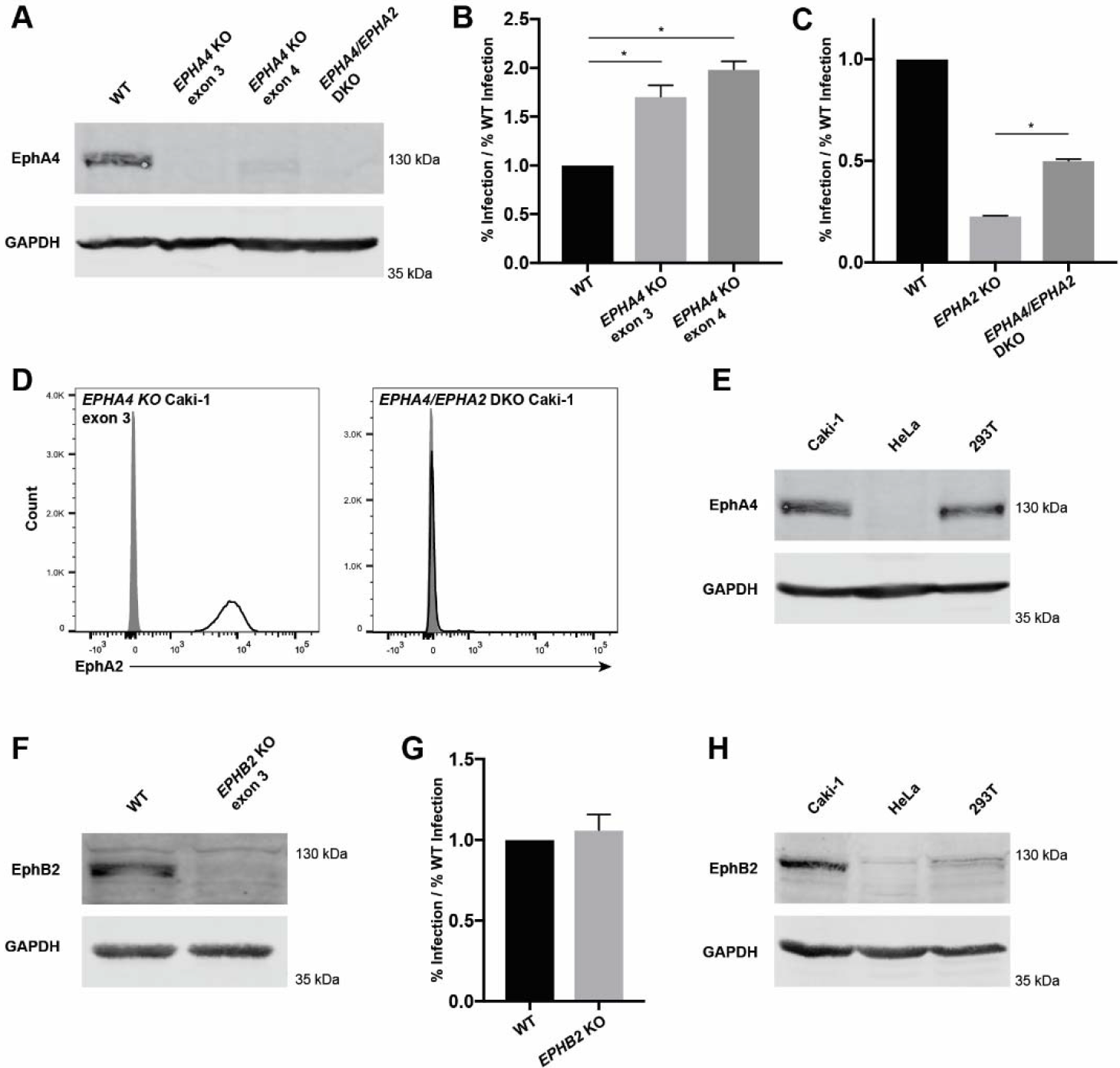
EphA4 and EphB2 are dispensable for infection of Caki-1 cells. (A, E, H) 120 μg or (F) 50 ug of the indicated whole cell lysate proteins were run on a 10% SDS-PAGE gel and blotted for EphA4 (A, E) or EphB2 (F, H) and GAPDH as a loading control. WT and *EPHA4* KO Caki-1 cells (B), or WT, *EPHA2* KO, and *EPHA4/EPHA2* DKO Caki-1 cells (C) were infected with KSHV in triplicate and infection percentage was measured by flow cytometry two days post infection. The infection percentages of KO cell lines were normalized to the average infection percent of WT cells and a representative experiment is shown. (D) *EPHA4* KO and *EPHA4/EPHA2* DKO Caki-1 cells were immunostained for surface EphA2 expression. Grey histograms represent isotype controls. (G) WT and *EPHB2* KO Caki-1 cells were infected with KSHV in triplicate and infection percentages were quantified by flow cytometry two days post infection. The infection rates of the KO line were normalized to the average WT infection rate and a representative experiment is shown. *, p < 0.05.

WT Caki-1 cells were transfected with px330 plasmids containing guide sequences targeting *EPHA4* and *EPHB2* (Table 1). We were unable to find an antibody that reliably detected EphA4 or EphB2 by surface immunostaining of live cells, so single-cell clones were derived from the transfected populations and then screened for the loss of EphA4 or EphB2 by western blot. Two *EPHA4* KO Caki-1 cell lines and one *EPHB2* KO Caki-1 cell line were isolated (Figs. 5A, 5F). WT, *EPHA4* KO, and *EPHB2* KO Caki-1 cell lines were then infected with KSHV. Surprisingly, the infection percentage of *EPHA4* KO cells was elevated compared to WT cells, while the infection percentage of *EPHB2* KO cells was not significantly different (Figs. 5B, 5G). Similar results were observed in *EPHA4* and *EPHB2* KO Caki-1 cells created with a lentiviral CRISPR-Cas9 system (data not shown).

To further understand the infection phenotype of *EPHA4* KO cells, one *EPHA4* KO Caki-1 cells (exon 3) were transfected with the *EPHA2*-targeted px330 plasmid (Table 1) and a pool of *EPHA2/EPHA4* DKO cells was isolated by FACS (Figs. 5A, 5D). When these cells were infected with KSHV, the infection percentage was reduced compared to WT cells, but significantly elevated compared to *EPHA2* KO cells (Fig. 5C). These data suggest that either EphA4 is a negative regulator of KSHV infection, or that Caki-1 cells compensate for the loss of EphA4 in a way that enhances KSHV infection.

EphA4 and EphB2 were detected in Caki-1 lysate and EphA4 was additionally found in 293T lysate. Importantly, both of these proteins were lacking in HeLa cell lysate, even though EphA2-independent infection was observed in both Caki-1 and HeLa cells (Figs. 5E, 5H). Furthermore, these results show that EphA4 and EphB2 are dispensable for KSHV infection and are unlikely to be the functional targets of ephrin-A4-Fc blocking during infection of *EPHA2* KO Caki-1 cells.

### Multiple Eph receptors rescue KSHV infection of EphA2 KO cells

Although we found that two endogenous Eph receptors besides EphA2 were not required for KSHV infection in WT Caki-1 cells, the significant infection defect of *EPHA2* KO Caki-1 cells provided an ideal platform to test the effects of transduced Eph receptors on KSHV infection. The clonal *EPHA2* KO Caki-1 cell line, described above, was used for these experiments.

To ensure that expression levels of different Eph receptors could be compared, mature forms of *EPHA2, EPHA4*, and *EPHA5* lacking endogenous signal peptides were cloned into p3xFlag-CMV-9 following the preprotrypsin leader sequence and a 3xFlag tag (Fig. 6A). This cloning scheme ensured that the proteins would be properly oriented in the membrane during translation and ultimately be N-terminally tagged with 3xFlag. The 3xFlag-tagged Eph receptor constructs were cloned into a retroviral vector and transduced into *EPHA2* KO Caki-1 cells.

**Figure 6.**
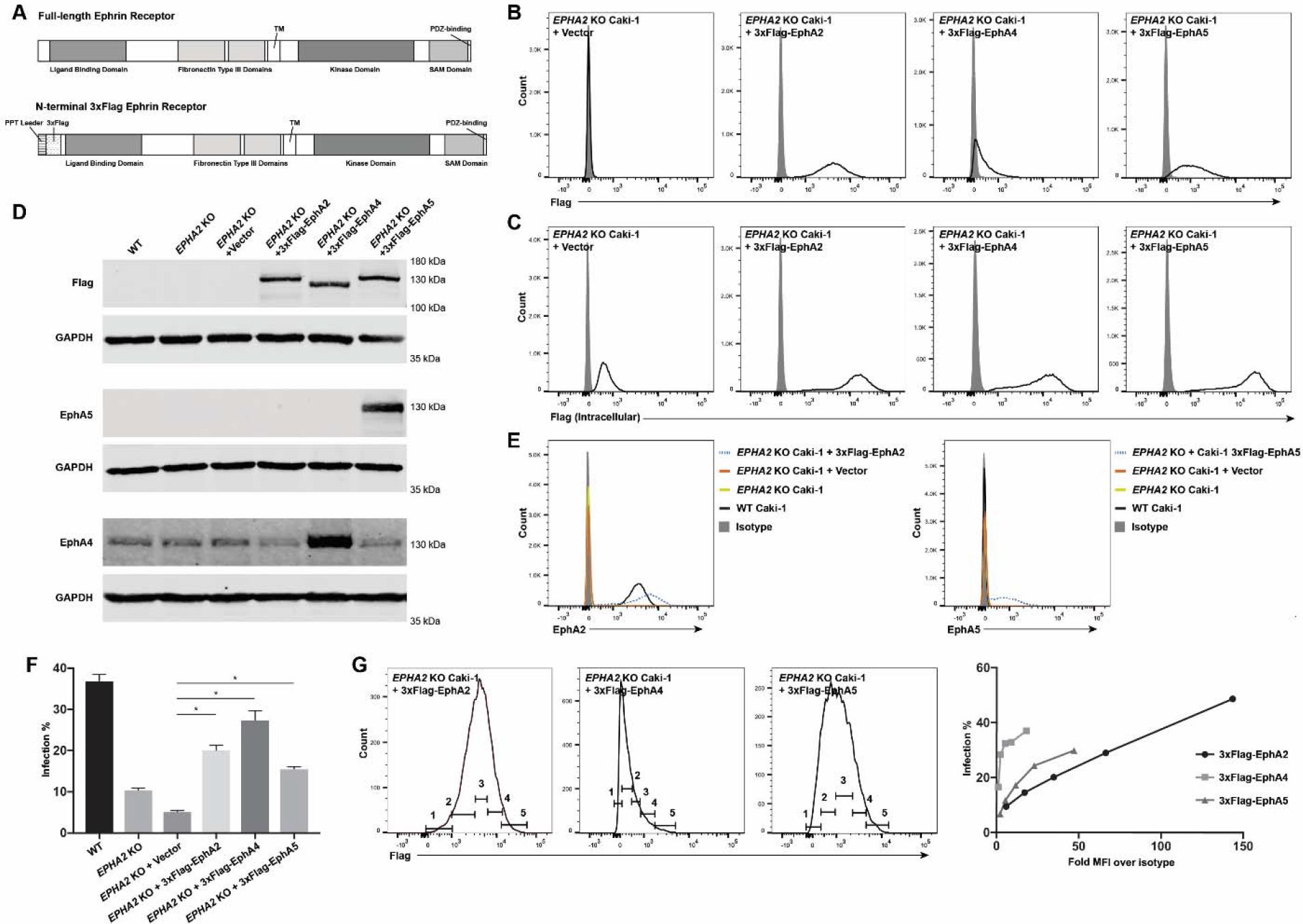
EphA2, EphA4, and EphA5 rescue KSHV infection in *EPHA2* KO Caki-1 cells. (A) Diagram of generalized full-length and PPT-3xFlag-mature ephrin receptor constructs. Live (B) or fixed and permeabilized (C) 3xFlag-tagged ephrin receptor transduced *EPHA2* KO cells and a vector control were immunostained for surface (B) or intracellular (C) 3xFlag expression and analyzed by flow cytometry. Grey histograms represent isotype controls. (D) The indicated cell lysates were run on 10% SDS-PAGE gels and blotted for 3xFlag, EphA4, and EphA5 with matched GAPDH as a loading control. For the Flag and EphA5 blots, 15 μg of whole cell lysate protein was loaded. For the EphA4 blot, 120 μg of whole cell lysate protein was loaded. (E) The indicated cell lines were immunostained for surface EphA2 or EphA5 expression and analyzed by flow cytometry. Grey histograms represent isotype controls. (F) The indicated cell lines were infected with KSHV in triplicate and infection rate was quantified by flow cytometry two days post infection. A representative experiment is shown. (G) The 3xFlag expression histograms of infected 3xFlag-tagged ephrin receptor transduced cell lines were divided into five successive gates as shown. The infection rate within each gate was plotted against the fold MFI over isotype of each gate. *, p < 0.05.

The 3xFlag tag was detected on the surface of each cell line by flow cytometry, although the magnitude of expression varied with each receptor (Fig. 6B). However, when the cell lines were stained for intracellular 3xFlag, the overall expression levels of the receptors appeared to be similar to each other (Fig. 6C). Additionally, when the expression of the receptors was examined by western blot, the intensities of the 3x-Flag-tagged bands were similar across the three transduced cell lines (Fig. 6D). These data show that all three constructs were expressed to a similar degree but that cell surface trafficking of the three receptors was different.

The expression of the 3x-Flag-tagged Eph receptors was also compared to the corresponding endogenous protein. The peak of surface expression of transduced 3xFlag-EphA2 was slightly higher than endogenous EphA2 as measured by flow cytometry, but the range of EphA2 expression in the population of 3xFlag-EphA2-transduced cells was much greater compared to WT cells (Fig. 6E). EphA5 was not naturally expressed by Caki-1 cells, but EphA5 was readily detected in 3xFlag-EphA5-transduced cells as a wide peak by flow cytometry and also as a strong band by western blot (Figs. 6E, 6D). In contrast, EphA4 was found to be expressed endogenously by Caki-1 cells by western blot and the EphA4 band became more pronounced in the 3xFlag-EphA4-transduced cell lysate (Fig. 6D).

WT Caki-1, *EPHA2* KO Caki-1, and the 3xFlag-tagged Eph receptor transduced *EPHA2* KO cell lines and a vector control were infected with KSHV. The surface 3xFlag expression was measured concurrently with infection percentage by flow cytometry. 3xFlag-EphA2, 3xFlag-EphA4, and 3xFlag-EphA5 all significantly rescued the infection percentage to varying degrees compared to the vector control (Fig. 6F). Because of the broad range of flag-tagged receptor expression within the populations of the transduced cell lines, we also examined how the infection percentage changed with surface protein level. The histograms of 3xFlag expression from one replicate well of the experiment were divided into five successive gates (Fig. 6G). The percent GFP+ cells in each bin was plotted against the fold geometric mean of the fluorescence intensity (MFI) over isotype MFI of each bin (Fig. 6G). Surprisingly, we found that both EphA4 and EphA5 promoted higher KSHV infection rates at lower amounts of surface protein compared to EphA2. However, at very high surface expression levels that were only attained by EphA2, the infection percentage surpassed that of WT cells from the same experiment (Fig. 6F).

Altogether these data show that EphA2, EphA4 and EphA5 can rescue the infection rate phenotype of *EPHA2* KO cells which suggests that the function of EphA2 in this infection mechanism may not be specific to EphA2. Moreover, the overexpression of EphA4 in this context strongly enhanced KSHV infection, which is not what we expected based on our *EPHA4* KO experiments. The precise role of endogenous EphA4 cannot be discerned from these studies alone.

### Ectodomain of EphA2 is sufficient to rescue KSHV infection in EphA2 KO cells

In primary microvascular endothelial cells and HFFs, several studies have reported that downstream effector proteins co-immunoprecipitate and colocalize with EphA2 during infection, implying involvement of the cytoplasmic tail of EphA2 which contains a kinase and several protein-protein interaction domains (40–43, 48). EphA2 is also phosphorylated upon KSHV infection in HEK293 and SLK cells (39, 48) Here we have shown that KSHV infection of Caki-1 and HeLa cells is independent of canonical KSHV integrin receptors, but still dependent on EphA2. Eph receptors can naturally trigger endocytosis in response to ephrin ligand binding by several mechanisms (reviewed in 60). Since KSHV gH/gL binds to the ephrin-binding domain of EphA2 (44, 49), we hypothesized that the signaling domains in the cytoplasmic tail of EphA2 would be necessary for infection and would provide clues about how the virus might use EphA2 to enter cells without utilizing canonical integrin receptors.

To this end, a truncation mutant of EphA2 was generated which lacked all cytoplasmic signaling domains. This cytoplasmic truncation (ΔCT) contained the entire peptide signal and ectodomain (amino acids 1-537) and the transmembrane (TM) domain (aa 538-558) (Fig. 7A). Full length *EPHA2* (FL) and *EPHA2* ΔCT were cloned into retroviral vectors and stably transduced into *EPHA2* KO Caki-1 and HeLa cells. Both EphA2 FL and EphA2 ΔCT were expressed on the cell surface to a slightly higher degree than endogenous EphA2 on Caki-1 (Fig. 7D) and HeLa cells (data not shown). When infected with KSHV, both EphA2 FL and EphA2 ΔCT significantly rescued infection to nearly identical levels compared to the vector control (Figs. 7B, 7C).

**Figure 7.**
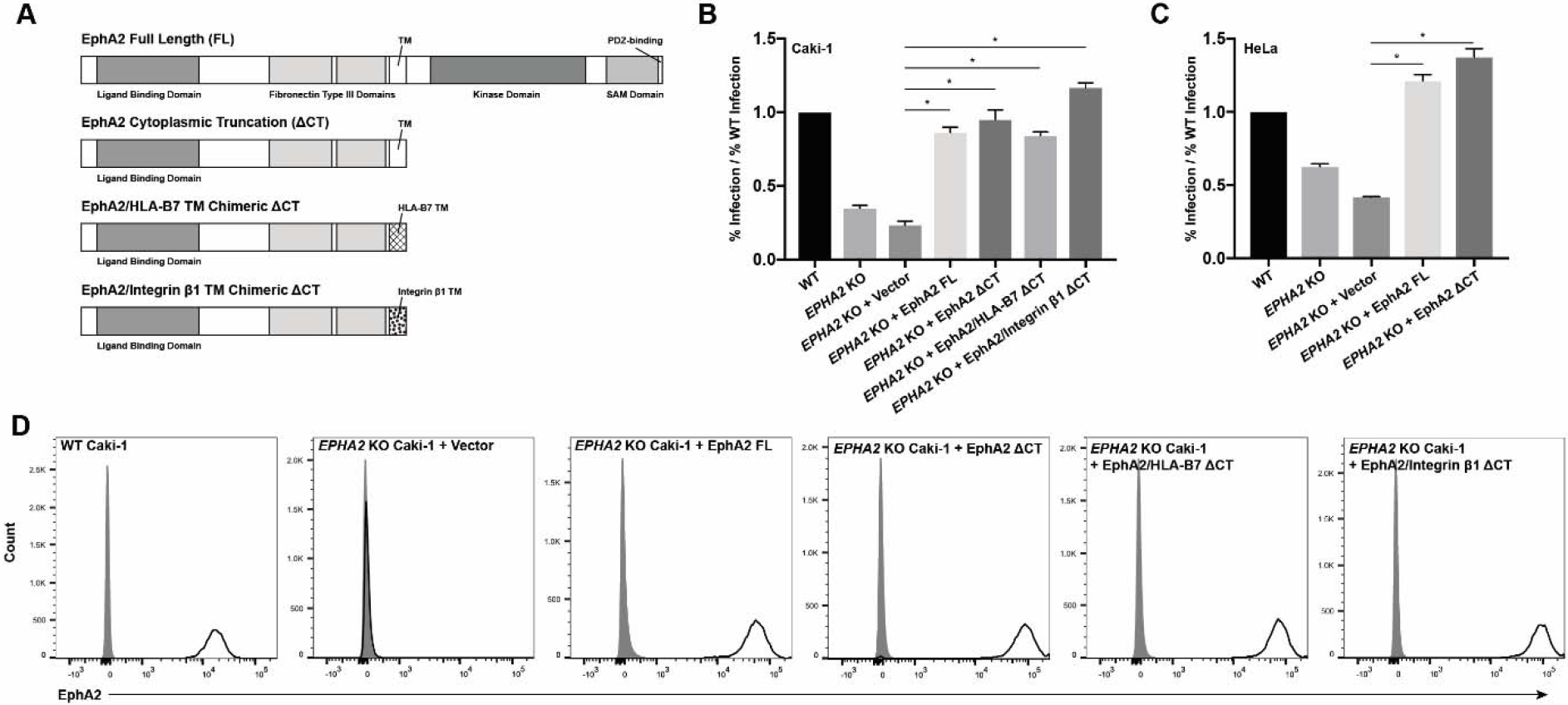
EphA2 ectodomain is sufficient to rescue infection rate in *EPHA2* KO cells. (A) Diagram of EphA2 truncation and domain swap constructs. (B, C) WT, *EPHA2* KO, and *EPHA2* KO cells transduced with the EphA2 constructs indicated in (A) were infected with KSHV in triplicate and infection rate was quantified by flow cytometry two days post infection. The infection rates were normalized to the average infection rate of WT cells and a representative experiment is shown. (D) WT, *EPHA2* KO, and the indicated transduced *EPHA2* KO Caki-1 cells were immunostained for surface EphA2 expression and analyzed by flow cytometry. Grey histograms represent the isotype controls. *, p < 0.05.

Eph receptor clustering is essential for natural signaling events, and a homodimerization region has been found within the transmembrane domain of EphA1 by nuclear magnetic resonance spectroscopy (61). To test whether the transmembrane domain was required for EphA2 ectodomain function during KSHV infection, we performed additional domain swaps with the EphA2 ΔCT construct and replaced the EphA2 TM domain with that of integrin β1 or HLA-B7, two unrelated single-pass transmembrane proteins (Fig. 7A). These domain-swapped constructs were also cloned into retroviral vectors and transduced into *EPHA2* KO Caki-1 cells. The EphA2/HLA-B7 chimeric CT and EphA2/integrin β1 chimeric ΔCT constructs were expressed at the cell surface to the same degree as the FL and CT constructs, and they also significantly rescued KSHV infection of *EPHA2* KO cells (Figs. 7D, 7B). Together, these data show that only the ectodomain of EphA2 is required to rescue KSHV infection in *EPHA2* KO Caki-1 cells.

### Infection of primary gingival keratinocytes requires HS interactions

Since transmission through saliva is thought to be a major route of KSHV infection, we examined the expression and use of known KSHV receptors during KSHV infection of primary gingival keratinocytes (PGKs). First, surface expression of known KSHV receptors on PGKs was examined by flow cytometry. We found that HS, EphA2, and the integrin subunits α3, αV, β1, and β5 were readily detected at the cell surface (Fig. 8A). Like HeLa cells, PGKs did not express integrin β3 at the cell surface, nor did we detect xCT or DC-SIGN (Fig. 8A).

**Figure 8.**
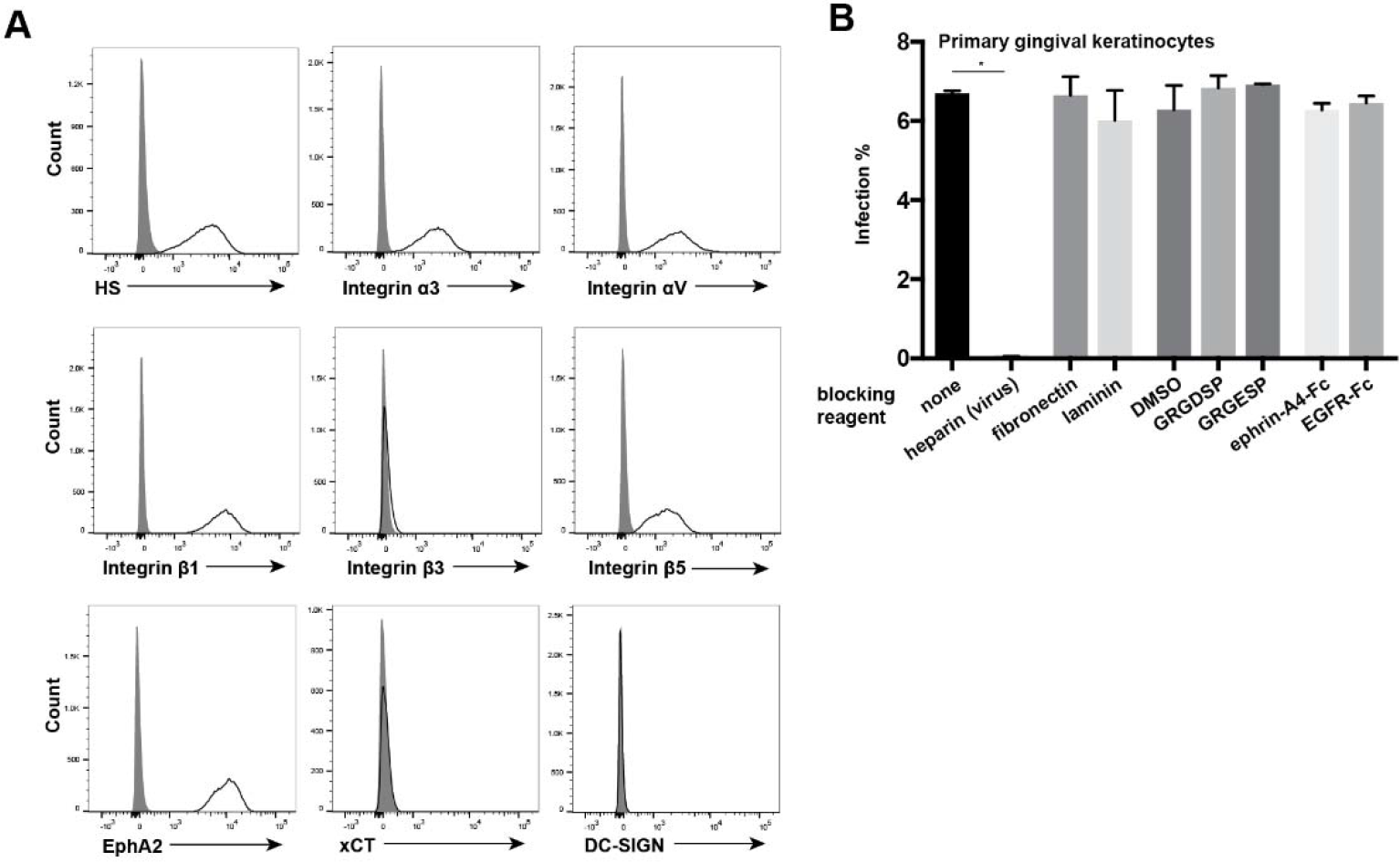
KSHV infection of PGKs depends on HS interactions but is not inhibited by integrin- or Eph-blocking agents. (A) PGKs were immunostained for surface expression of known KSHV receptors and analyzed by flow cytometry. Grey histograms represent isotype controls. (B) PGK cells were preincubated with fibronectin or laminin at 50 μg/L, GRGDSP or GRGESP peptides at 2 mM or an appropriate volume control of DMSO, and ephrin-A4-Fc or EGFR-Fc as a control at 5 μg/mL for one hour at 4°C. For the no treatment and heparin condition, cells were pre-incubated in normal media at 4°C. For the heparin block condition, virus was blocked with heparin at 500 μg/mL for one hour at 37°C. Cells were then washed and infected in triplicate with KSHV, or heparin-blocked KSHV for two hours at 37°C. Ephrin-A4-Fc and EGFR-Fc concentrations were maintained during the infection. Infection percentage was quantified by flow cytometry two days post infection. *, p < 0.05.

Next, we utilized the blocking experiments that we had replicated from existing KSHV receptor literature to test whether these known receptors were utilized during infection of PGKs. To test whether HS interactions were required for infection, KSHV was pre-blocked with heparin before infection. To investigate whether any canonical integrin receptors were required for infection, cells were pre-blocked with fibronectin, laminin, or the RGD-containing peptide GRGDSP, the control peptide GRGESP, or a volume control of DMSO. Finally, to investigate whether Eph receptor interactions were required for infection, cells were pre-blocked with ephrin-A4-Fc or EGFR-Fc as a control. The cells were then washed and infected with KSHV (or heparin-blocked KSHV in the HS test condition) for two hours.

We found that heparin block of virions completely abrogated the infection of PGKs, as was the case for WT HeLa cells (Figs. 8B, 2D). Similar to our results with Caki-1 and HeLa cells, the integrin ligands and RGD peptide had no significant effects on infection percentage in PGKs (Fig. 2.16). Surprisingly, the ephrin-A4 ligand also had no effect on infection in these cells (Fig. 2.16). These results suggest that KSHV infection of PGKs does not require interactions with the laminin-binding integrin α3β1, the RGD-binding integrin αVβ5, nor EphA2 which is competitively blocked by ephrin-A4 (39, 47). However, infection of PGKs clearly requires heparan sulfate interactions.

## DISCUSSION

In this report, we describe a novel KSHV infection mechanism in Caki-1 and HeLa cells which requires HS and the ectodomain of EphA2 but is independent of the canonical KSHV integrin receptors α3β1, αVβ3, and αVβ5. We also present evidence that infection of primary gingival keratinocytes (PGKs) is dependent on HS but not EphA2 or the canonical KSHV integrin receptors. Finally, we found that EphA4 and EphA5 may regulate KSHV infection in various contexts. CRISPR-Cas9 proved to be a valuable tool to dissect the roles of individual receptors during KSHV infection.

It is thought that HS broadly acts as an attachment factor for many viruses including KSHV, but some publications indicate that HS may have additional functions during KSHV infection of several cell types. One study reported that HS was required on target HEK293, CHO, and human conjunctival epithelial cells in a virus-free fusion assay with effector cells that expressed KSHV gB, gH, and gL, suggesting that HS is involved in the interactions between KSHV glycoproteins and entry receptors (46). A second study used advanced imaging to reveal that upon initial binding to HT1080 fibrosarcoma cells KSHV only colocalized with HS about half of the time, while colocalization with canonical integrin receptors was much more robust (35). However, soluble heparin still inhibited KSHV binding to these cells (62). Our experiments clearly demonstrated that HS interactions are required for KSHV to infect Caki-1, HeLa, and PGKs, but the precise role of HS during infection remains an open question. Interestingly, the blocking effect of heparin appeared to be more severe on HeLa cells and PGKs which both lacked surface integrin β3. A possible explanation for this is that integrin αVβ3 could play a role in attachment during the infection of Caki-1 cells, especially in the *EXT1* KO context. Likewise, our integrin subunit KO studies were performed in the context of HS-expressing cells, which could possibly mask a functionally redundant role shared by HS and certain integrin heterodimers.

Surprisingly, we found that KSHV infection was unaffected by perturbations in the integrin network in Caki-1 and HeLa cells despite the well-characterized roles that integrins α3β1, αVβ3, and αVβ5 play during infection of HFF and primary endothelial cells (reviewed in 11). However, these results are in agreement with several studies in which integrin-blocking reagents failed to inhibit KSHV infection of HEK293 and SLK cells (32, 39, 50). We found that Caki-1 and HeLa cells lacking either integrin αV or β1— abolishing the expression of five and twelve integrin heterodimers, respectively—were infected to a similar percentage as WT cells. We also found no infection defect in Caki-1 cells knocked out for both integrin subunits α3 and αV, effectively lacking integrins α3β1, αVβ3, and αVβ5.Furthermore, a panel of integrin ligands and RGD peptides had little or no effect on the percent of Caki-1 cells, HeLa cells, or PGKs infected by KSHV.

Our CRISPR-Cas9 KO studies covered sixteen of the twenty-four known integrin heterodimers, and we further determined that the remaining eight heterodimers were not expressed in Caki-1 cells. It is still conceivable that an αV-family and one or more β1-family integrins besides α3β1 are fully redundant receptors of KSHV in this system, although such a situation would not be consistent with several past studies where a KSHV infection phenotype was recorded after targeting only a single integrin heterodimer with a blocking antibody (34, 36, 45).

It should be noted that the results of our experiments with HeLa cells may not be in agreement with a recent KSHV receptor study on a HeLa-derivative cell line that was misidentified as human salivary gland epithelial cells (HSG[HeLa]) (35, and corrected in 52). This study reported that HSG(HeLa) cells were mostly resistant to infection, despite expressing HS, EphA2, xCT, and integrins α3β1 and αVβ5 (35, 52). Like our HeLa CCL-2 cells, HSG(HeLa) cells did not express integrin αVβ3. The infection rate of HSG(HeLa) cells was greatly increased upon expression of integrin β3 leading the authors to conclude that integrin αVβ3 was a crucial receptor for KSHV in these cells (35, 52). The differing conclusions from this study and ours may be attributed to the experimental approaches used, since our work focused on depleting receptors from WT cells instead of overexpressing them. It is also possible that HSG(HeLa) cells and our HeLa CCL-2 cells are too divergent to be comparable, as it is unclear how far removed HSG(HeLa) cells are from parental HeLa strains.

The KSHV glycoprotein gB binds integrins through an RGD domain that mimics natural integrin ligands, as well as a DLD domain (34, 36, 50, 59). In HFF and primary microvascular endothelial cells, this gB-integrin interaction is required to initiate the KSHV-induced signaling cascade through the activation of focal adhesion kinase and other downstream effectors that eventually lead to virion endocytosis (36–38). This leads to the outstanding question of how KSHV internalizes in Caki-1 and HeLa cells without the involvement of canonical integrin receptors. We hypothesized that KSHV might directly induce endocytosis through EphA2, mimicking natural ephrin ligand-receptor binding events. Several studies report phosphorylation of EphA2 during KSHV infection and suggest that the cytoplasmic domain of EphA2 is essential to propagate KSHV-induced signaling events and recruit effectors of macropinocytosis and clathrin-mediated endocytosis, but this idea has never been directly tested in the context of *EPHA2* KO cells (39–43, 48). While we found that infection of Caki-1 and HeLa cells required EphA2, remarkably an EphA2 construct truncated after the TM domain rescued infection in *EPHA2* KO cells as efficiently as the full-length EphA2 construct. Furthermore, infection of PGK cells was unaffected by competitive blocking with ephrin-A4, which has been previously shown to efficiently inhibit infection in multiple cell types (39, 47–49). It is possible that in certain cellular contexts, EphA2-mediated signaling is not necessary for infection but could be activated or recruit effector proteins as a bystander effect.

Together, our results suggest that in Caki-1 cells, HeLa cells, and PGKs, KSHV does not trigger the same integrin-EphA2 signaling axis that is so crucial for entry into HFF and primary microvascular endothelial cells. However, this conclusion must be rectified with the strong infection defect we observed in *EPHA2* KO Caki-1 and HeLa cells. The KSHV membrane glycoprotein gH/gL binds strongly to EphA2 (39, 44, 47, 49), so one interpretation of our data is simply that the ectodomain of EphA2 acts as an adhesion receptor in the cellular context of Caki-1 and Hela cells. EphA2 could also have this function in PGKs, but functional redundancy with another protein may mask its role.

Taken together with our experiments investigating potential roles for different Eph receptors during KSHV infection, more speculative hypotheses can also be made. The result that we were able to further inhibit infection of *EPHA2* KO cells with ephrin-A4 suggests that another factor which is blocked by ephrin-A4, most likely an Eph receptor, promotes KSHV infection. We ruled out that endogenous EphA4 and EphB2 were necessary for infection of Caki-1 infection, but also found that these Eph receptors were not expressed by HeLa cells. This is important because we found that HeLa cells also exhibit a significant amount of EphA2-independent KSHV infection, even more than Caki-1 cells. It is possible that additional Eph receptors are expressed by both cell lines and affect KSHV infection in *EPHA2* KO and WT contexts.

In support of this idea, we demonstrated that transduced EphA4 and EphA5 constructs rescued infection rates in *EPHA2* KO cells to levels comparable with transduced EphA2. In fact, at low amounts of surface expression, EphA4 and EphA5 constructs outperformed EphA2 in this assay. It is unclear why endogenous EphA4 is dispensable for and perhaps even inhibits infection in the endogenous setting, while it promoted KSHV infection in the overexpression context. Spliced or modified forms of EphA4 produced from the endogenous gene could account for this discrepancy. Alternatively, EphA4 may be part of a homeostatic network that ultimately impacts KSHV infection efficiency and cellular adaptation to the loss of EphA4 could be responsible for the KO phenotype. Whether EphA4, EphA5, and other Ephs act as true cellular receptors for KSHV or otherwise regulate KSHV infection should be further investigated.

A striking property of Eph receptors is that they form heterotetramers with their ligands as well as large oligomerized arrays through Eph-Eph interactions in their ectodomains which are critical to trigger forward signaling in response to ligands (63–66). These signaling arrays can contain multiple types of Eph receptors, Eph receptors that are not bound to ligands, and Eph receptor ectodomains (63, 64, 66–68). Moreover, Eph cluster size, composition, and the presence of alternatively spliced Eph receptor forms may all influence the cellular outcomes of Eph signaling (69–71). Importantly, one study showed that the ectodomain of EphA2 was sufficient to localize the protein to cell-cell contacts (63), and another study of chimeric EphA2 and EphA4 constructs suggested that the ectodomain may be a stronger determinant of cellular responses than the attached cytoplasmic domain (65). Thus, it is conceivable that in the presence of other signaling-competent Eph receptors, the ectodomain of EphA2 could promote clustering and signaling during KSHV infection just as well as the full-length receptor as we observed in our experiments.

However, it is also possible that an unknown factor―not an Eph receptor―is responsible for virion internalization and EphA2-independent infection in Caki-1 and HeLa cells. In support of this, a new study has identified the motif in gH of KSHV and rhesus rhadinovirus (RRV) that is required for Eph receptor binding (49). When this motif was mutated, *de novo* KSHV infection of HFF and endothelial cells was severely attenuated at the post-attachment stage, but not completely blocked (49). Additionally, infection of HFF, endothelial cells, and SLK/Caki-1 cells with this mutant could no longer be blocked with soluble forms of EphA2 or ephrin-A4 (49). Not only is this study consistent with our *EPHA2* KO data in Caki-1 and HeLa cells, the existence of another KSHV receptor may explain why infection of PGK cells was not inhibited by soluble ephrin-A4. This unknown receptor hypothesis is not exclusive to the potential involvement of other Eph receptors. It is possible that Eph receptors regulate each other’s activity in modulating another KSHV receptor in certain cellular contexts.

It is still unclear why targeting EphA2 with either CRISPR-Cas9 or ephrin-A4 had such differential effects on Caki-1 and HeLa cells versus PGKs. Eph receptor signaling is known to be quite cell type-dependent, and therefore the availability of EphA2 in the cell membrane or its intracellular signaling outcomes may naturally differ in PGKs. However, EphA2 has also been found to be upregulated in many types of solid tumors and its intracellular signaling functions may also be dysregulated in this context, possibly complicating the interpretation of our results in the Caki-1 and HeLa cell lines (reviewed in 72 and 73).

Interestingly, two independent groups recently discovered that EphA2 is a receptor for the gammaherpesvirus Epstein-Barr Virus (EBV) on epithelial cells (74, 75). While integrins αVβ5, αVβ6, and αVβδ had previously been identified as epithelial cell receptors for EBV (76, 77), one group demonstrated with CRISPR-Cas9 KO cells that αV-family integrins were not required for EBV glycoprotein-mediated fusion with HEK293 cells (74). Furthermore, these studies demonstrated that the kinase activity of EphA2 and indeed the entire intracellular domain were dispensable for EBV glycoprotein fusion and infection, respectively (74, 75). These results are strikingly similar to the findings we report here, and further characterization of both infection mechanisms may reveal more parallels in receptor use between these two related gammaherpesviruses.

Given the importance of epithelial cell infection for host colonization, it will be valuable to further characterize KSHV entry in the absence of canonical integrin receptors and its impact on the viral life cycle. Our data support the notion that KSHV receptor usage and entry mechanisms vary widely between cell types. We propose that KSHV infection is not restricted by integrin and EphA2 expression and that the virus may utilize several members of both the integrin and Eph receptor families in various combinations for entry into a broad variety of cell types throughout the body. Modern gene editing technologies such as CRISPR-Cas9 will facilitate detailed studies of KSHV receptors in the future and have the potential to rapidly expand the field of virus-host interactions.

## MATERIALS AND METHODS

### Cell lines and culture

SLK/Caki-1 (ATCC HTB-46) cells were a gift from D. Ganem. HeLa cells (ATCC CCL-2) were obtained from the UC Berkeley BDS Cell Culture Facility. HEK293T cells (ATCC CRL-1573), Phoenix cells (ATCC CRL-3213), and primary gingival keratinocytes (ATCC PCS-200-014) were purchased from the ATCC. Primary gingival keratinocytes were grown in Dermal Cell Basal Medium (ATCC PCS-200-030) supplemented with Keratinocyte Growth Kit (ATCC PCS-200-040) at 37°C with 5% CO_2_. All other cells were grown in high glucose Dulbecco’s Modified Eagle’s Medium (DMEM, Gibco) supplemented with 5% fetal bovine serum (FBS, SeraDigm) at 37°C with 5% CO_2_.

### KSHV production and infection

iSLK.Bac16 (gift from J. Jung, USC, see 78) cells harboring latent KSHV.BAC16 infection were cultured under selection with 1.2 mg/mL of hygromycin B (Invitrogen). The cells were induced to produce virus with 1 mM sodium butyrate (Alfa Aesar) and 1 ug/mL doxycycline (Sigma-Aldrich). Three days after reactivation, supernatant was collected and filtered through a 0.45 μm syringe filter. The unconcentrated viral supernatant was stored at 4°C and diluted with standard culture medium for use in infection experiments. The dilution was calculated for each batch for an infection percentage of ~30% on WT Caki-1 cells after 24hrs of infection, measured in GFP+ events by flow cytometry. Cells were incubated with virus for 12-24 hours, then viral supernatant was removed and replaced with fresh medium until analysis two days post infection. For blocking experiments, cells were incubated with virus for only two hours, then washed and grown in fresh medium.

### CRISPR-Casθ genome editing

Guide sequences were designed using the online tool crispr.mit.edu and are provided in Table 1. A 5’ G was added to sequences that didn’t already contain one and then the appropriate adaptors were appended to both forward and reverse oligos to facilitate cloning into px330 (Addgene #42230) according to the protocol provided at genome-engineering.org. Assembled px330 plasmids were transfected into cells of interest and mutant cells were sorted by FACS or subcloned to obtain genetic KO cell pools or cell lines, respectively.

### Antibodies

Heparan sulfate antibody (F58-10E4) was purchased from Amsbio, integrin α3 antibody (P1B5 from Calbiochem, integrin αV, integrin β7, EphA2, and EphA5 antibodies (MAB12191, MAB4669, AF3035, and MAB541, respectively) from R…D Systems, integrin β1 and integrin β3 antibodies (T2S/16 and PM6/13, respectively) from Novus Biologicals, integrin β5 and EphA2 antibodies (AST-3T and SHM16, respectively) from BioLegend, xct and GAPDH antibodies (ab37185 and ab181602, respectively) from Abcam, DC-SIGN antibody (DCN47.5) from Miltenyi Biotec, EphA4 antibody (4C8H5) from ThermoFisher, EphB2 antibody (2D12C6) from Santa Cruz Biotech, and Flag antibody (M2) from Sigma-Aldrich. Purified isotype control antibodies (MAB002, MAB003, MAB004, AB-105-C, MAB006) were purchased from R…D Systems except mouse IgM, κ (MM-30) was from BioLegend.

### Blocking reagents

Ephrin-A4-FC and EGFR-Fc were purchased from R…D Systems. GRGDSP and GRGESP peptides were purchased from Anaspec. Human fibronectin and mouse laminin were purchased from Corning.

### Constructs and cloning

Eph receptors were amplified from BJAB (EphA4, EphA5) or Caki-1 (EphA2) cDNA and directly cloned into pQCXIN (Clontech) or cloned into p3xFlag-CMV-9 (Sigma-Aldrich) and subsequently cloned into pQCXIN to add an N-terminal 3xFlag tag preceded by the preprotrypsin leader sequence. Truncation mutants were amplified with a reverse primer in the indicated position containing an artificial stop codon. Chimeric TM domain EphA2 constructs were made using SOEing PCR.

### Transfection and transduction

Caki-1 and HeLa cells were transfected with px330 and phoenix cells were transfected with pQCXIN-based constructs using Fugene transfection reagent (Promega) and Optimem (Gibco) according to the manufacturer’s instructions. After 2-3 days, retrovirus was collected from the phoenix cell supernatant and filtered through a 0.45 μm filter. Filtered retroviral supernatant was applied to target cells with 6 μg/mL polybrene (Santa Cruz) and spinfected at 500xg for 2 hours at room temperature. Transduced cells were selected with neomycin (Fisher Scientific) at 1.2 mg/mL.

### Flow cytometry and sorting

Cells were harvested with trypsin (Gibco) or PBS (Gibco) + 2 μM EDTA (Fisher) when staining for trypsin-sensitive epitopes. Cells were blocked, stained, and washed in 1% BSA (Fisher) in PBS. When applicable, cells were fixed in 4% PFA (ThermoFisher Pierce) in PBS and permeabilized with 0.25% Triton X-100 (EM Science) in PBS. Live cells were stained with DAPI (BioLegend) and fixed cells with Ghost Dye Violet 510 (Tonbo Biosciences) for viability according to the manufacturer’s instructions. Cells were analyzed using an LSR Fortessa or LSR Fortessa X-20 cell analyzer (BD Biosciences) and sorted using a BD Influx or BD FACSARIA Fusion cell sorter (BD Biosciences). Data was processed and visualized with FlowJo 10 (BD Biosciences).

### Western blotting

Cells were harvested by scraping in PBS and lysed in RIPA buffer (150 mM sodium chloride, 1% Triton X-100, 0.5% sodium deoxycholate, 0.1% sodium dodecyl sulfate, 50 mM Tris, and a protease inhibitor cocktail (Roche)). Protein concentration in lysate was quantified by BCA assay (ThermoFisher Pierce). Lysates were run on a 10% polyacrylamide gel and transferred to a nitrocellulose membrane. A buffer containing 3% BSA and 10% FBS in TBST (20 mM Tris, 150 mM sodium chloride, 0.1% Tween 20) was used for blocking and primary antibody incubation. Plain TBST was used for washing and secondary antibody incubation. Blots were visualized with IRDye 800CW and 680RD secondary antibodies from LI-COR Biosciences using a LI-COR Odyssey infrared scanner and analyzed in ImageStudio Lite 5.2 (LI-COR Biosciences).

### Statistical analysis

The indicated data sets were compared using the student’s t-test in Prism 7 (GraphPad). A p value < 0.05 was denoted with a ^*^.

## ACKNOWLEDGEMENTS

This research received no specific grant from any funding agency in the public, commercial, or not-for-profit sectors. The authors would like to thank Hector Nolla and Alma Valeros from the University of California, Berkeley Flow Cytometry Core Facility for technical assistance and advice, and Ella Hartenian and Dr. Britt Glaunsinger for helpful data analysis which helped guide the experiments. The authors would like to additionally thank Dr. Ellen Robey, Dr. Eva Harris, Dr. Trever Greene, Andrew Birnberg, Kristina Geiger, and Valerie Vargas-Zapata for enlightening discussion about this project.

